# Folding of Aquaporin 1: Multiple evidence that helix 3 can shift out of the membrane core

**DOI:** 10.1101/014167

**Authors:** Minttu T. Virkki, Nitin Agrawal, Elin Edsbäcker, Susana Cristobal, Arne Elofsson, Anni Kauko

**Affiliations:** Department of Biochemistry and Biophysics Science for Life Laboratory, Stockholm University SE-171 21 Solna, Sweden; Department of Biosciences, Biochemistry, Åbo Akademi, FI-20520 Turku, Finland; Department of Clinical and Experimental Medicine, Cell Biology, Faculty of Health Science, Linköping University, Linköping, Sweden and Departments of Physiology, IKERBASQUE, Basque Foundation for Science, Faculty of Medicine and Dentistry, University of the Basque Country, Leioa, Spain; Department of Biochemistry and Biophysics and Science for Life Laboratory, Stockholm University SE-171 21 Solna, Sweden

**Keywords:** Membrane protein, Translocon recognition, Protein folding, Hydrophobicity, Molecular dynamics

## Abstract

The folding of most integral membrane proteins follows a two-step process: Initially, individual transmembrane helices are inserted into the membrane by the Sec translocon. Thereafter, these helices fold to shape the final conformation of the protein. However, for some proteins, including Aquaporin 1 (AQP1), the folding appears to follow a more complicated path. AQP1 has been reported to first insert as a four-helical intermediate, where helix 2 and 4 are not inserted into the membrane. In a second step this intermediate is folded into a six-helical topology. During this process, the orientation of the third helix is inverted. Here, we propose a mechanism for how this reorientation could be initiated: First, helix 3 slides out from the membrane core resulting in that the preceding loop enters the membrane. The final conformation could then be formed as helix 2, 3 and 4 are inserted into the membrane and the reentrant regions come together. We find support for the first step in this process by showing that the loop preceding helix 3 can insert into the membrane. Further, hydrophobicity curves, experimentally measured insertion efficiencies and MD-simulations suggest that the barrier between these two hydrophobic regions is relatively low, supporting the idea that helix 3 can slide out of the membrane core, initiating the rearrangement process

## Introduction

α-helical integral membrane proteins are essential for signaling, transport, energy production and catalysis. The majority of α-helical membrane proteins fold following a two-stage process^1^. First, sufficiently hydrophobic segments are inserted into the membrane by the Sec translocon^2,3^ thereafter the protein folds. When a segment is sufficiently hydrophobic it is recognized by the Sec-translocon^2, 4, 5^ and the orientation of the segment is primarily guided by the preference of positively charged residues in cytosolic loops^6^. The initial recognition is followed by the less well-studied assembly of transmembrane segments, binding of co-factors and formation of reentrant regions^7^.

Although the folding process is less well understood recent studies have highlighted that the folding process can in fact be remarkably complicated. In Bacteriorhodopsin the folding transition state is not reached until the second and last (seventh) helix interact^8^. Further, addition of positively charged residues can completely flip EmrE^9^ as well as most of the N-terminal half of LacY changes orientation in response to altered lipid composition^10^. In Cystic Fibrosis Transmembrane Conductance Regulator the integration of transmembrane helices proceeds in an unexpected order, the first transmembrane helix is integrated into the membrane only after the second transmembrane helix has been inserted^11^. These observations indicate that hydrophilic regions might pass through the membrane during folding.

The cost associated with hydrophilic regions passing through the membrane during these large-scale rearrangements could potentially be overcome by the utilization of external machineries, such as the translocon^12^. However, the membrane is not a uniform hydrophobic slab, but rather dynamic, and at least partly permitting the passage of polar groups. Indicative of this is how charged cell penetrating peptides^13,14^, pore forming peptides^15^ and some C-tail anchored proteins^16,17^ can enter the membrane spontaneously. The ability of polar groups to draw lipid head-groups and water deep into the core may provide a mechanism for their entry to the membrane^18,19^.

A particular illustrative example for large-scale rearrangements is Aquaporin 1 (AQP1). Aquaporins forms water-soluble pores in biomembranes and in addition to the six transmembrane helices they contain two reentrant regions^20^. These two regions come together to almost form a seventh helix. Antibody epitope experiments in *Xenopus oocytes* demonstrated that AQP1 insert initially as a four-helix intermediate and only later folds into its final structure^21^. In this intermediate, helices 2 and 4 are not inserted in the membrane, and consequently helix 3 is inserted in an inverted orientation^21^, see Figure 1. Based on experiments in mammalian cells the existence of the four-helix intermediate was initially questioned^22^, but this contradiction has been explained by the observation that the intermediate is less stable in mammalian cells^23^. In contrast the close homolog Aquaporin 4 (AQP4) follows the conventional folding pathway, where each transmembrane segment is co-translationally inserted into the membrane^21^.

**Figure 1:**
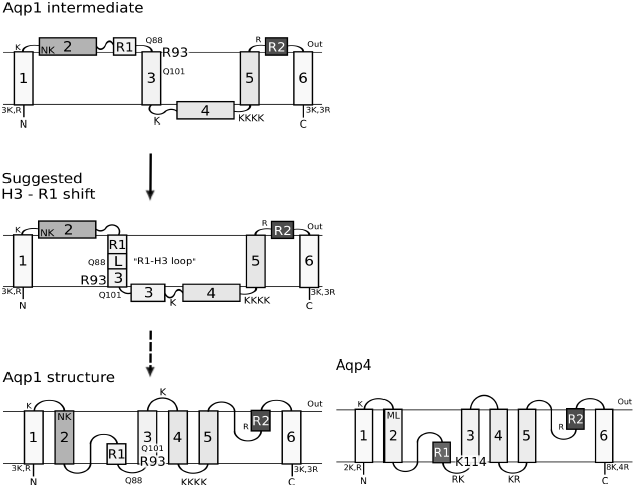
An overview of the topologies of AQP1 and AQP4 and the proposed “R1-H3 shift”. The proposed folding pathway for AQP1 is shown next to the topology of AQP4. All positively charged residues are shown. The grey shading roughly depicts the hydrophobicity of each segment to highlight the differences in hydrophobicity of TM2 and R1 between AQP1 and AQP4. The location of Asn49 and Lys51 in AQP1 helix 2 and the corresponding Met70 and Leu72 in AQP4 are also depicted. These residues are responsible for the hydrophobicity differences between the second helices^24^. AQP1 is initially inserted into the membrane as a four-helix intermediate and later folds into its final six-helix topology^21^. This requires the reorientation of helix 3. Here, we propose that helix 3 may spontaneously shift out of the membrane core (the R1-H3 shift), initiating the folding, despite the presence of polar residues (Arg93, Glu88, Glu101). Helix 3 in AQP4 contains three positively charged residues at its N-terminal side causing and orientational preference not present in the corresponding positions in AQP1.

In order to understand the sequence features causing the differences in folding pathways between AQP1 and AQP4 both proteins have been studied by truncation-reporter experiments in dog pancreatic microsomes^24^. The most notable differences are; Helix 2 in AQP1 is less hydrophobic as it contains two polar residues, Asn49 and Lys51^24^. When AQP1 helix 2 is not integrated into the membrane helix 3 is inserted in an inverted orientation, see Figure 1. The positive inside effect then prevents the integration of helix 4, as the C-terminal loop contains four lysines^22^. In addition, the loops flanking helix 3 have been suggested to play a role for its orientation^24^.

In this study we aim to shed some light on the folding process of AQP1. Based on our earlier observation that large-scale shifts are not infrequent in helical membrane proteins^25^, we propose that helix 3 can shift out of the membrane core and bring the preceding R1-H3 loop into the membrane, see Figure 1. We propose that this “R1-H3 shift” serves as a first step in AQP1 folding, followed later by additional events. Using a combination of experimental and computational techniques we find that the “R1-H3 shift” is a feasible first step in AQP1 folding.

## Results and Discussion

Here, we show that the “R1-H3 shift” is feasible and could serve as a first step in the folding of AQP1. In this model the third transmembrane helix of AQP1 shifts out of the membrane core and the preceding “R1-H3 loop” is brought into the membrane. We show that the R1-H3 loop is sufficiently hydrophobic to reside in the membrane and that the energetic cost of the shift is consistent with the model. In contrast, the corresponding regions in AQP4 do not have these characteristics.

### The R1-H3 loop is more hydrophobic in AQP1 than in AQP4

The hydrophobicity profiles of AQP1 and AQP4 protein families show the conservation of hydrophobicity profiles within each family but differs between the two families, see Figure 2. In AQP1 helix 2 is less hydrophobic and a hydrophobic segment (ΔG_pred_≈ 0 kcal/mol) is present just before helix 3. In AQP4 this segment is less hydrophobic (ΔG_pred_≈ 4 kcal/mol). This hydrophobic segment contains the helical section of reentrant region 1, the loop between the reentrant region and helix 3, and the N-terminal part of helix 3. Below we will refer to this region as the “R1-H3 loop”. The hydrophobicity of the R1-H3 loop indicates that this region might insert into the membrane. The hydrophobic barrier between helix 3 and the R1-H3 loop is relatively low (ΔG_pred_≈3 kcal/mol) and is mainly caused by a single residue, Arg93.

**Figure 2:**
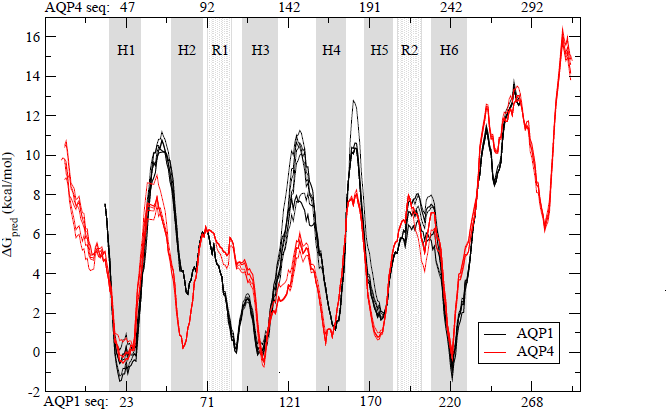
Hydrophobicity plots for aligned AQP1 and AQP4 protein sequences. Transmembrane helices are shaded in in dark grey and reentrant regions in light grey. Sequence numbering corresponding to *Homo sapiens* AQP1 is shown below the figure and the numbering corresponding to AQP4 from *Rattus norvegicus* is shown on top. The predicted free energy of insertion (ΔG_pred_) for each residue was calculated using the Hessa scale^5^. The hydrophobicity profiles are conserved in both families, but show a clear difference between the families. In AQP1 the R1-H3 loop is almost as hydrophobic as helix 3 and the barrier separating these regions is quite low.

During the “R1-H3 shift” Arg93 has to cross the membrane from the lumenal side to the cytosolic side, see Figure 1. Further, the positive inside rule would favor the rearrangement and support the “R1-H3 shift” hypothesis, which would serve as the first step in AQP1 folding.

### In AQP1 the R1-H3 loop can be inserted into the membrane

A well-established *in vitro* glycosylation assay was used to identify segments in AQP1 and AQP4 that can be recognized by the translocon^5,26^. Glycosylation can only occur in the microsome lumen and can therefore be used as a topology marker. The insertion efficiencies of potential transmembrane helices can be determined by separating constructs with different number of attached glycans on SDS-PAGE, see Figure 3.

**Figure 3:**
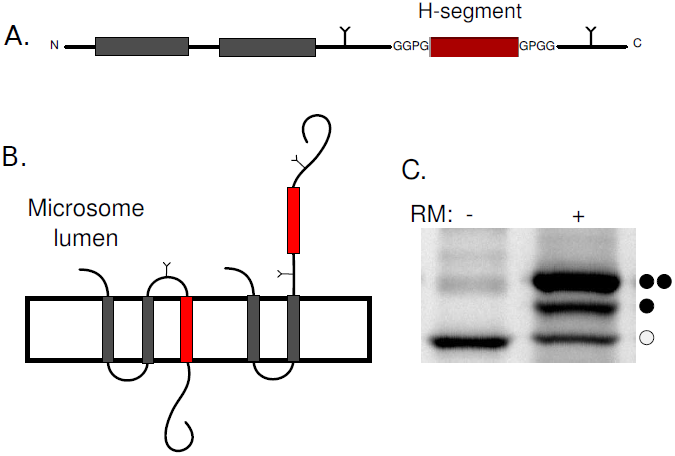
The leader peptidase (Lep) as a host protein and the *in vitro* expression in the presence of microsomes system. A) Segments from AQP1 and AQP4 were introduced as H-segments (depicted here in red) into the P2 domain of Lep, preceded by the two transmembrane helices (shown in grey) of wild type Lep. Lep is known to insert in a N_1um_ – C_1um_ orientation in rough microsomes. Asn-Xaa-Thr glycosylation sites were introduced on both sides of the H-segment. B) As glycosylation by the oligosaccharyl transferase only occurs in the microsome lumen, the topology of a construct can be deduced from the number of glycans added to the protein. Here, singly glycosylated species arise from proteins with the H-segment inserted into the membrane whereas double glycosylated species arise from a translocated H-segment. C) An example of in vitro translation in the absence (-) and presence (+) of microsomes. Each glycan adds around 2 kDa to the molecular mass of the protein allowing their separation on SDS-PAGE. Here, non-glycosylated, single and double glycosylated protein species are indicated with open circles, one filled and two filled circles respectively. In this example about 75% of the protein is doubly glycosylated.

Insertion efficiencies of the R1-H3 loop (Ala78-Ala100), helix 3 (Ala94-Thr116) and the least hydrophobic segment between them (Leu84-Ile106) in AQP1 and corresponding segments from AQP4 were tested. In Figure 4 the experimental (ΔG_exp_) values are shown together with the calculated hydrophobicity values (ΔG_pred_). Given the experimental limitations the predicted and experimental values are in good agreement. In AQP1 both the R1-H3 loop and helix 3 insert well, while in AQP4 only helix 3 is recognized by the translocon.

**Figure 4:**
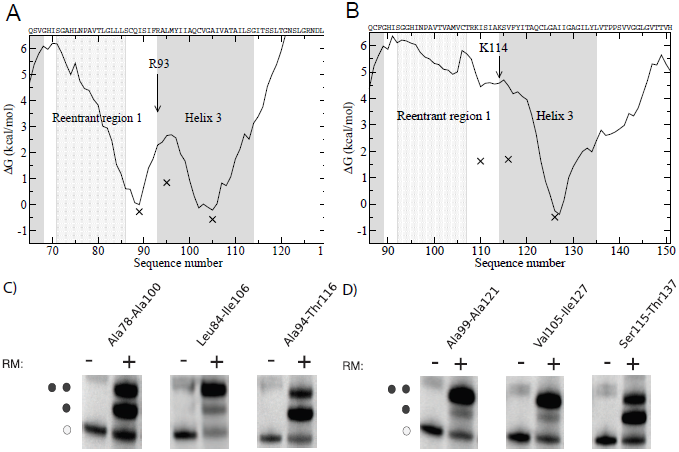
Experimentally determined insertion efficiencies compared to calculated insertion efficiencies. ΔG_exp_ values are plotted against the central position of the 23 residues long peptide. The curves represent the hydrophobicity as measured by ΔG_pred_. Helices are shown with grey and reentrant region with a light grey background. A) In AQP1 both helix 3 and the R1-H3 loop area can be recognized by the translocon as independent membrane segments. In addition, the barrier is relatively low, which supports the possibility of a shift. B) In AQP4 only helix 3 is efficiently recognized as a transmembrane segment by the translocon. C) Representative SDS-page gels for AQP1 constructs expressed *in vitro*. D) Representative SDS-page gels for AQP4 constructs expressed *in vitro*.

Translocon recognition was also tested using longer segments. These all start before the R1-H3 loop and include various truncations of helix 3. In general these long segments inserts slightly better than the shorter segments, see Table 1. In AQP1, but not in AQP4, the segment truncated at Ala100 that only includes six residues of helix 3 is inserted well, i.e. the R1-H3 loop is efficiently recognized by the translocon in AQP1.

**Table 1:** Insertion efficiencies, experimentally measured and predicted free energies for segments from AQP1 and AQP4 The central position of the most central residue in the most hydrophobic segment is shown in column 3.

| Construct | Length | Hydrophobic center | Inserted | $\Delta G_{\text{exp}}$ | $\Delta G_{\text{pred}}$ |
| --- | --- | --- | --- | --- | --- |
| *Aquaporin 1* |  |  |  |  |  |
| His69-Met96 | 28 | 89 | 45% | 0.12 | 0.85 |
| His69-Ala100 | 32 | 91 | 80% | -0.83 | -0.05 |
| Ala78-Ala100 | 23 | 91 | 61% | -0.26 | 0.01 |
| Ser86-Tyr97 | 11 | 92 | 6% | 1.7 | 4.9 |
| Leu84-Ile106 | 23 | 96 | 18% | 0.92 | 2.70 |
| Ala94-Val103 | 11 | 100 | 7% | 1.6 | 4.5 |
| Ala94-Thr116 | 23 | 106 | 72% | -0.56 | -0.20 |
| His69-Gly125 | 57 | 107 | 87% | -1.14 | -0.20 |
| *Aquaporin 4* |  |  |  |  |  |
| His90-Ala113 | 24 | 105 | 15% | 1.04 | 3.80 |
| His90-Phe117 | 28 | 105 | 15% | 1.04 | 3.80 |
| His90-Ala121 | 32 | 111 | 17% | 0.95 | 3.80 |
| Ala99-Ala121 | 23 | 111 | 6% | 1.68 | 4.45 |
| Val105-Ile127 | 23 | 117 | 5% | 1.69 | 4.48 |
| Ser115-Thr137 | 23 | 127 | 69% | -0.48 | -0.30 |
| His90-Thr137 | 60 | 127 | 80% | -0.83 | -0.53 |

### Identifying the translocon-recognized segments

Next, we aimed to determine the exact boundaries for the translocon-recognized segments in AQP1 using Minimal Glycosylation Distance Mapping (MGD)^27^, see Figure 5. Two different constructs were studied: His69-Gly125, which contains the R1-H3 loop and the full-length helix 3, and His69-Ala100, which only contains six residues from helix 3.

**Figure 5:**
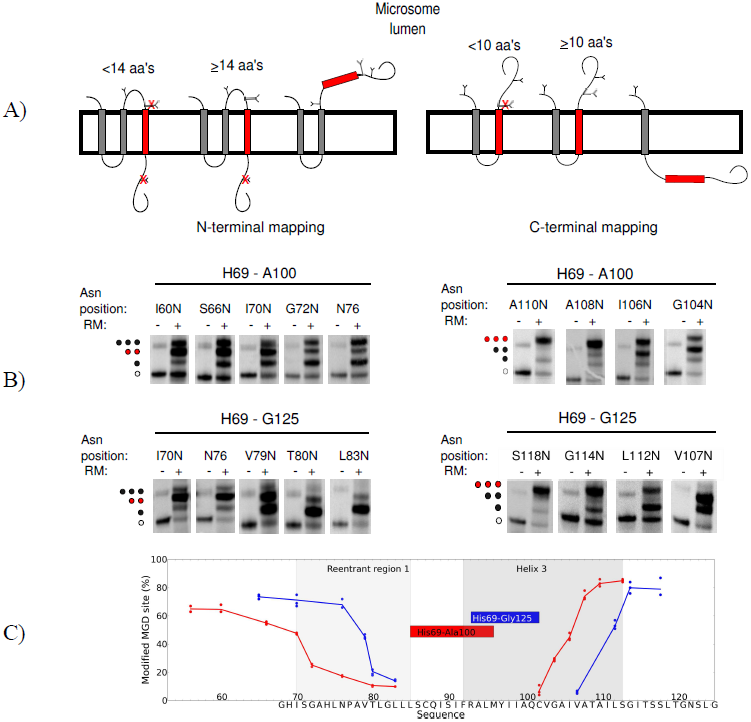
Minimal glycosylation distance mapping (MGD) of the N- and C-terminal ends of transmembrane domains of His69-Gly125 and His69-Ala100 from AQP1. A) N-terminal mapping is shown to the left and C-terminal mapping to the right. In MGD, a third glycosylation site (MGD-site) is placed either before or after the H-segment. The position of the MGD-site is placed at different distances from the transmembrane region. The entire construct is expressed *in vitro* and the fraction of glycosylation at the MGD-site is measured. When approximately 50% of the MGD-sites are modified, that position is known to reside ≈ 14 residues before the N-termini of the membrane embedded region and ≈ 10 residues of its C-termini. B) The SDS-page gels show MGD-mapping for the two constructs. The position of the MGD-sites are indicated on top of the gels with the position corresponding to where the H-segments are embedded into the membrane marked in red. C) The glycosylation efficiency of the MGD-site is plotted against its position. The R1 and H3 regions are marked with light and dark grey shading, while the red and blue boxes represent the transmembrane helices as determined by the MGD-mapping for His69-Ala100 and His69-Gly125 respectively. The sequence of His69-Gly125 is presented at the X-axis.

In the longer construct the glycosylation mapping identifies Ala94 to be the first membrane embedded residue and Val103 to be the last, see Figure 5. The N-terminus is in perfect agreement with the membrane boundary found in the crystal structure of AQP1 as defined in the PDBTM database^28^, but the helix is truncated at its C-terminus. In the shorter construct the identified membrane region is shifted towards the N-terminus, spanning residues Ser86 to Tyr97. The C-terminus is located at the position expected for the recognition of the R1-H3 loop, but also this helix appears shorter than what is expected for a transmembrane helix.

One possible explanation for these unexpectedly short transmembrane regions could be that the segments are located in multiple membrane locations. They may be able to slide like a piston from one side of the membrane to the other. Then, when performing MGD mapping, the glycosylation could kinetically trap a state shifted towards one of the sides. Thereby, when mapping the C-terminus it would be shifted, and the helix appear to be shorter. The identified short membrane regions are not recognized by the translocon, see Table 1. This demonstrates, that while the termini of these segments can reside within the membrane one at a time, they cannot simultaneously be inside the membrane. Hence, a piston like motion seems likely to occur, and when the glycosylation site is modified, the peptide is trapped.

### Understanding the mechanisms enabling the R1-H3 shift

According to our experiments, the hydrophobic barrier between reentrant R1-H3 loop and H3 in AQP1 is ≈ 0.9*kcal/mol*, see Figure 4 and Table 1. This low barrier supports the possibility of a spontaneous R1-H3 shift, which involves Arg93 crossing the membrane. To obtain insights into the molecular details of this transition, a series of molecular dynamics simulations were performed where the AQP1 R1-H3 peptide was restrained to different positions in the membrane, see Figure 6.

**Figure 6:**
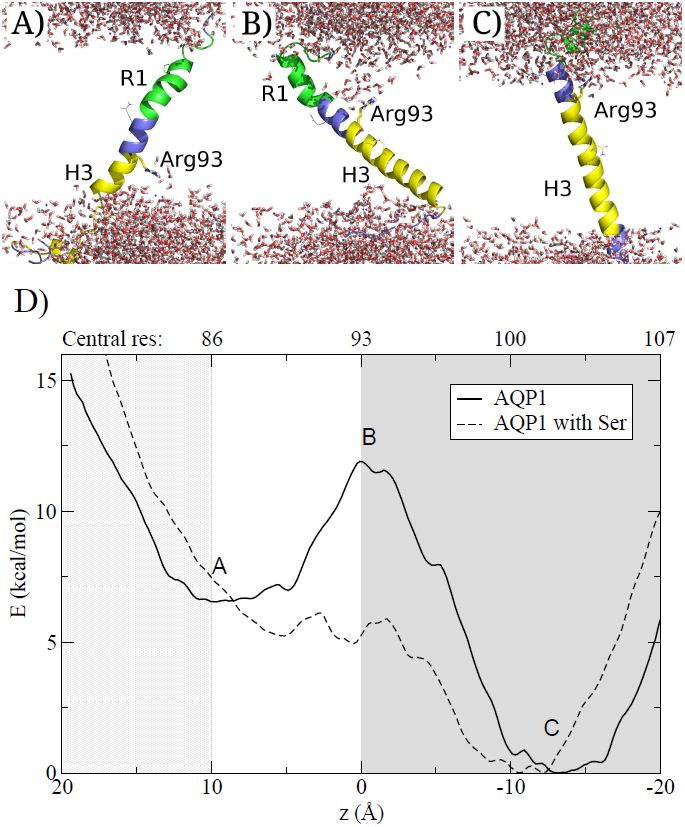
A-C) Snapshots from the simulation of the AQP1 R1-H3 segments with Arg93 placed at different positions in the membrane. Arg93 is depicted as a stick model. Gln88, Gln101 and Pro77 (that caps the reentrant region helix) are shown as lines. The helix 3 is drawn in yellow, loops in blue and R1 in green. Water molecules are depicted as stick models and no lipid molecules are shown for clarity. A) R1-H3 region in the membrane, B) Arg93 in the middle of the membrane C) Helix 3located in the middle of the membrane. D) Calculation of the potential mean force from umbrella samplings of the AQP1 R1-H3 segment. The mean force curves show two minima corresponding to when helix 3 or R1-H3 loop are in the membrane. The position where the snapshots are taken are marked with letters. The barrier corresponds Arg93 at the membrane. When serine analogs are added to the membrane, the barrier is decreased to near biological level. The letters A-C refer to the structural images. The number corresponding to the residue that located in the center of the membrane is shown above the plot. H3 is shown in grey and R1 in light grey.

When either the R1-H3 loop or helix 3 are located at the membrane center, the long side-chain of Arg93 snorkels towards one of the membrane interfaces. However, when Arg93 is in the center of the membrane, the membrane becomes distorted and water enters the membrane, as has been seen in earlier studies^29,30^. Further, to avoid the energetic cost of dissolving hydrophobic regions into the surrounding water, the peptide tilts and brings the hydrophobic segment of helix 3 into the membrane. This might further lower the cost of the barrier.

Further, potential of mean force (PMF) calculations were used to estimate the free energy cost of insertion from the simulations. As expected for AQP1, the profile contains two clear minima, corresponding to the R1-H3 loop and helix three, Figure 6D. The barrier between these minima corresponds to when Arg93 is in the center of the membrane.

The general shape of the PMF curve is similar to what is observed experimentally but the energy barriers are higher. This is a well-known phenomena^31^ and in accordance with earlier studies^3, 29^. The discrepancy has been explained by the lowered polarity of the translocon interior^32,31^ or by increased polarity caused by membrane proteins^33,34^. To approximate for polar residues within the membrane, simulations were also performed with twenty serine analogs located in the membrane. In this system the barrier is lowered, indicating that the R1-H3 shift may be plausible in real cytoplasmic membranes.

### Implications for folding of AQP1

The results presented above suggest a spontaneous shift of helix 3 during AQP1 folding. When helix 3 moves out of the membrane core the R1-H3 loop integrates into the membrane. The positive inside rule would favor such a shift, as Arg93 would move to the cytoplasmic side. Also Arg93 may form stabilizing interactions with other residues in AQP1 during the shift, in particular with Glu17 from helix 1. The reinsertion of helix 3 together with helix 4 would then be sufficient to bring helix 3 into its correct orientation, see Figure 1. How the hydrophilic H3-H4 loop could pass through the membrane is not clear. However, AQP1 is a channel and the reorientation process requires the presence of helices 5 and 6^21^. In addition, the final topology would require refolding of helix 2 and the reentrant regions. It could be imagined that these regions enter the membrane jointly.

The reorientation of AQP1 is less efficient in a cell free system than in *Xenopus oocytes* indicating that while the R1-H3 shift could occur spontaneously, the later stages of rearrangements may depend on the presence of additional cellular machineries^21^. The translocon has been suggested to play a key role in the reorientation of AQP1^12^. Alternatively, the translocating chain associated membrane protein (TRAM), could also be involved, as it has previously been shown to aid the insertion of charged helices^35^. On the other hand, LacY has been shown to be capable of dramatic reorientation, including change in orientation for transmembrane domains and post-translational insertion of a transmembrane helix, without any other cellular factors except the lipid composition of the membrane^36^. Anyhow, extensive additional studies are required to understand how the folding could proceed after the R1-H3 shift.

## Conclusions

The folding of AQP1 does not follow the traditional two-stage folding process. In AQP1 helix 3 inverts its orientation in the membrane after the initial insertion whereas this does not occur in the homologous AQP4. Consequently, AQP4 does not show any of the characteristics listed below^21^. Here, we propose a mechanism for the initial steps in the folding of AQP1; First helix 3 is shifted out of the membrane core resulting in the preceding regions to be pulled into the membrane, this is followed by a reinsertion of helix 3 in its correct orientation, see Figure 1. We present three observations supporting this idea. First, we noted an additional conserved hydrophobic segment, the R1-H3 loop, next to helix 3, see Figure 2. We show that this region can be integrated efficiently into the membrane by the translocon, see Table 1 and Figure 4. Secondly, experimental, predicted and simulated hydrophobicity values implicate a relatively low barrier between helix 3 and the R1-H3 loop, see Figure 2, 4 and 6. Also experimental minimum glycosylation distance mapping suggests that several alternative segments can be recognized by the translocon in the R1-H3 region and that this region might undergo a piston like motion. Finally, the positive inside rule would also favor the shift as Arg93 would move to the cytoplasmic side of the membrane, see Figure 1.

## Methods

### Alignments and ΔG plots

All members of the Aquaporin 1 and 4 families were extracted from Swissprot^37^ in Nov 2011. A multiple sequence alignment of all these protein sequences was done using kalign^38^ with default parameters. The hydrophobicity of individual segments was estimated by the predicted free energy of insertion (ΔG_pred_) calculated from the biological hydrophobicity scale^5^. For each residue in the hydrophobicity profiles the optimal window length ranging between 19 and 23 residues was used.

### Enzymes and chemicals

Unless otherwise stated, chemicals were obtained from Sigma-Aldrich (St. Louis, MO, USA), oligonucleotides were obtained from MWG Biotech AG (Ebersberg, Germany) and all enzymes were from Fermentas (Burlington, Ontario, Canada), except Phusion DNA polymerase that was obtained from Finnzymes OY (Espoo, Finland). The plasmid pGEM-1 and the TNT® SP6 Quick Coupled Transcription/Translation System were from Promega Biotech AB (Madison, WI). [^35^S]Met was bought from Perkin Elmer (Boston, MA) and the column washed dog pancreas rough microsomes were from tRNAprobes (College Station, Texas). The EndoH assay kit was from New England Biolabs (Ipswich, MA). The Qiaprep Miniprep Plasmid Purification kits from QIAGEN (Hilden, Germany) were used for plasmid purifications. E.Z.N.A Cycle Pure and Gel Extraction kits from Omega Bio-Tek (Norcross, GA) were used during post-PCR manipulation.

### DNA manipulation

The *lepB* gene had previously been introduced into the pGEM-1 vector under the control of the SP6-promoter^39^ and with the context 5’ of the initiator codon changed to a Kozak consensus sequence^40^. To allow Lep to "host" other protein segments, SpeI and KpnI restriction recognition sites had been introduced in the sequence encoding the middle of the P2-domain (LepI). In all constructs two glycosylation sites are placed on each side of the H-segment^3^. To insulate each sequence from the Lep sequence, all segments contain both N- and C-terminal `GGPG` … `GPGG` flanks.

Double-stranded oligonucleotides encoding the different protein segments from AQP1 (*Homo sapiens)* and AQP4 (*Rattus norvegicus)* were introduced into the *lepB* gene as SpeI-KpnI-fragments of amplified PCR fragments using primers complementary to the 5’ and 3’ ends of the selected part of the gene (for exact amino acid sequences for these segments, see Table 1. Both the vector and the PCR fragments were digested with SpeI and KpnI (Fermentas) separated on agarose gel and fragments of correct size were excised from gel and purified (E.Z.N.A. Gel Extraction kit). PCR fragments were ligated to the vector carrying the *lepB* gene using Rapid DNA Ligation Kit (Fermentas).

### Point mutations

For minimal glycosylation distance mapping in AQP1 additional glycosylation sites were introduced into the sequence, using His69-Ala100 and His69-Gly125 as templates. To determine the first N-terminal residue recognized by the translocon, the H-segments were introduced into LepI and a series of constructs where the third glycosylation site was introduced at different positions downstream of Arg93 were made, see Figure 5A. For determining the last membrane embedded residue at the C-termini the constructs were introduced to LepII and again, a series of constructs with a third glycosylation site at varying distances upstream were made, Figure 5B.

For each glycosylation position the native amino acids in the sequence were exchanged into Asn-Ser-Thr. We chose to use the same glycosylation site sequon even if this introduces larger changes in the amino acid sequence of the H-segments to prevent fluctuations in glycosylation efficiency^41,42^. All DNA modifications were confirmed by sequencing of the plasmid DNA at Eurofins MWG Operon.

### Expression *in vitro*

Constructs in pGEM-1 were transcribed and translated in the TNT SP6 Quick Coupled System from Promega. A master mix containing [^35^S]-methionine (5 *μ*Ci) and lysate were mixed together in such a way that the amount of lysate is ten times the volume of [^35^S]-methionine. 5.5 *μ*l of this master mix was then added to 100 ng DNA. For positive reactions, the master mix was supplemented with dog pancreas column washed rough microsomes, in an amount that would yield at minimum 80 % targeting. All samples were incubated at 30°C for 90 min. For long segments the results did not differ more than 5 percentage points and therefore only doublets were made. However, as the variations were greater for AQP1 short segments, up to seven replicates were made. For MGD-mapping three replicates were made for each construct, with variation typically within 7 percentage points.

### Separation and analysis of expressed proteins

Translated proteins were separated by SDS-PAGE and visualized with a Fuji FLA-3000 phosphoimager (Fujifilm, Tokyo, Japan) with the ImageReader V1.8J/Image Gauge V 3.45 software (Fujifilm). The MultiGauge software was used to create one-dimensional intensity profiles for each lane on the gels, where the triply glycosylated proteins yield a higher molecular weight (+6 kDa), doubly glycosylated (+4 kDa) band and singly glycosylated proteins (+2 kDa), as compared to the non-glycosyalated protein. The differently glycosylated proteins are denoted with filled circles in the figures and the non-glycosylated protein with unfilled circles. Peak areas were then analyzed using the multi-Gaussian fit program from the Qti-plot software package (http://www.qtiplot.ro/).

### The apparent membrane insertion free energies for AQP1 and AQP4 segments

The apparent membrane insertion free energies ΔAG_ex_p for the H-segments (AQP1 and AQP4 segments given in Table 1) were calculated as follows. The fraction of singly (fx1) and doubly (fx2) glycosylated species can be used to calculate the apparent equilibrium constant 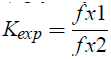 for a given H-segment. The *K*_*exp*_ value can be converted into an apparent free energy difference between the non-inserted and inserted state: Δ*G*_*exp*_ = –*RTlnK*_*exp*_, where R is the gas constant and T is the temperature in Kelvin. The accuracy of ΔG_exp_ determination is good between the interval of -1.5 and 1.0 kcal/mol^5^.

### Minimal glycosylation distance mapping

In MGD experiments the fraction of proteins that were singly, doubly and triply glycosylated was measured. Non-glycosylated proteins were not included, as they had not been targeted to the microsomal membranes.

For the N-terminal mapping, singly glycosylated proteins represent the state when the H-segment is integrated into the membrane and the MGD-site is too close to the membrane to be modified by the Oligosaccharyltransferase. For constructs that generate a larger fraction of doubly glycosylated proteins, the MGD-site is far enough from the membrane to get efficiently modified. H-segments that do not integrate into the membrane carry three glycans, see Figure 5. For the C-terminal mapping, doubly glycosylated proteins have the H-segment integrated into the membrane but the MGD-site is too close to the membrane. When the MGD-site is sufficiently distant a three-glycan form appear.

It should be noted that in the shorter construct the polar residues at the glycosylation site disturb the membrane insertion of the R1-H3 loop causing the short construct to not reach the same level of double glycosylation as in the longer construct, Figure 5. In addition, when the MGD-site is closer than 13 residues to Arg93, the segment is poorly recognized by the translocon as evident from gel images where a sudden increase of non-inserted (three-glycan form) can be seen, Figure 5C.

### Endoglycosidase H digestion

It was observed that some of the His69-Gly125 constructs in MGD-experiments appeared to be cleaved by Signal Peptidase resulting in multiple bands for some constructs, see Supplementary Figure S1. As the fraction of the constructs that are cleaved varies between differently glycosylated constructs it is necessary to identify which bands correspond to each glycosylation state.

In order to assess the correct cleaved fragments to their respective glycosylated protein species, an Endoglycosidase H (EndoH) assay was^26^ carried out. Here, 6 *μ*l of translation products were mixed with resuspended 1 *μ*l Denaturing Buffer (10x) and 3 *μ*l distilled water. After mixing, 2 ul of G5 Reaction Buffer (10x) and 7 *μ*l of distilled water were added. Finally, either 1 *μ*l of Endo H (500,000 units/ml) or dH2O (mock sample) was added. The samples were incubated at at 37 °C for 1 h. From here on, the samples were treated as all other *in vitro* expressed constructs. During analysis, in order to calculate the ratio of membrane embedded transmembrane regions the cleaved and non-cleaved forms were measured independently and the fractions non-cleaved and cleaved fragments were added.

### Molecular dynamics simulations

MD simulations were performed using Gromacs 4.5.5^43^ with the Berger lipid force field^44^. Equilibrated 1-Palmitoyl-2-oleoylphosphatidylcholine (POPC) and 1-Palmitoyl-2-oleoylphosphatidylserine (POPS) in a 3:1 mixture were used in the membrane bilayer. POPC was chosen, because it is widely used in MD simulations and its parameters are well optimized and POPS was included to represent anionic lipids that may interact with the cationic Arg93.

The length of AQP1 R1-H3 peptide was optimized by trial and error and the His69-Asn122 peptide was chosen. This peptide contains approximately an additional five residues on each side of the hydrophobic region. The peptide was built in an idealized helical conformation using the PyMOL Molecular Graphics System, Version 0.99, DeLano Scientific, Palo Alto, CA, USA. The peptide was embedded into a membrane by program g_membed^45^. The resulting membrane consisted of 66 POPC and 22 POPS molecules.

In all molecular dynamics simulations we used a 2 fs time step, LINCS constraints, 1.2 nm cutoffs (Coulomb, van der Waals and neighbor list), PME electrostatics, V-rescale temperature coupling to 323 K temperature and Parinello-Rahman pressure coupling to 1 bar pressure. A semi-isotropic temperature coupling was applied, where the pressure in the plain of the bilayer was coupled separately from the normal of the bilayer.

41 separate simulations were prepared with the R1-H3 segment positioned at different depth of the membrane. The starting position of each simulation differed by 0.1 nm. Each system was then solvated, neutralized by 21 sodium ions, minimized and equilibrated by a 1 ns simulation using position restraints (1000 *kJmol*^−1^nm^−2^) for all protein atoms. Each simulation was sampled during a 20 ns simulations using position geometry at the direction of the Z-axis and a 1000 *kJmol*^−1^*nm*^−2^ force constant. The membrane center and C-α atoms for residues 92-94 were used as reference groups. The first 5 ns of each simulation were discarded. Each simulation was run twice. To ensure full coverage of the mean force histogram, a few additional windows were added around the central position. The weighted histogram analysis method (program g_wham^46^) was used to extract the PMF curve.

It is well known that cellular membranes contain a large fraction of proteins. In simulations when additional proteins are included in the membrane, the cost for hydrophilic residues to enter the membrane is lowered^33^. However, in our simulations it was not possible to add entire helices, as this would have required much longer simulations, which would be computationally too expensive. Instead 20 serine analogs (methanol) were added to the membrane and each simulation was run three times. The addition of serines was based on the assumption that up to half of the membrane content consist of proteins, half of the residues are exposed and a quarter of exposed residues are at least mildly polar, i.e. 20 serine analogs roughly provides the same ratio of polar residues as found in real membranes^19^. The topology of the serine analog was modified starting from the topology of serine and the atom type of CB was changed to CH3. Serine was chosen, as it is a mildly hydrophilic residue that can act both as a donor and acceptor in hydrogen bond. Further it is rather frequent within membrane regions. Position restraints (1000 *kJmol*^−1^*nm*^−2^) were also applied to the Z-coordinate of the serine CB atom.

## Acknowledgments

Professor William Skach is kindly acknowledged for providing plasmids and harboring the genes for AQP1 and AQP4, Dr. Linnea Hedin and Dr. IngMarie Nilsson for valuable advice regarding laboratory work, Prof. Peter Tieleman for lipid structures, topologies and force field parameters, Prof. Erik Lindahl for stimulating discussions, Dr. Justin Lemkul for instructive web tutorials and Dr. Sara Light for proofreading the manuscript. AE, SC and MV were supported by grants from the Swedish Research Council, SSF, the Foundation for Strategic Research, Science for Life Laboratory the EU 6’th Framework Program is gratefully acknowledged for support to the GeneFun project, contract No: LSHG-CT-2004-503567 and the 7’th framework through the EDICT project, contract No: FP7-HEALTH-F4-2007-201924. SC was also supported by the Carl Trygger foundation. AK and NA were funded by the Finnish Academy and the Sigrid Juselius Foundation. Mark Johnson from Åbo Akademi University has provided to AK and NA excellent computing facilities (funded by Sigrid Juselius Foundation and Tor, Joe and Pentti Borg Foundation) and availability to the Biocenter Finland infrastructure and to the Finnish Grid Infrastructure. The majority of simulations were performed at the PDC center for high performance computing and we are also grateful for additional computational resources given to us by SNIC at NSC and Uppmax.

